# 53BP1 interacts with the RNA primer from Okazaki fragments to support their processing during unperturbed DNA replication

**DOI:** 10.1101/2023.09.27.559698

**Authors:** Melissa Leriche, Clara Bonnet, Jagannath Jana, Gita Chhetri, Sabrina Mennour, Sylvain Martineau, Vincent Pennaneach, Didier Busso, Xavier Veaute, Pascale Bertrand, Sarah Lambert, Kumar Somyajit, Patricia Uguen, Stéphan Vagner

**Author notes:** These authors contributed equally (Melissa Leriche, Clara Bonnet).

## Abstract

RNA-binding proteins are found at replication forks, but their direct interaction with DNA-embedded RNA species that inevitably shape physiological DNA replication remains unexplored. Here we report that 53BP1, involved in the DNA damage and replication stress response, is an RNA-binding protein that directly interacts with Okazaki fragments, in the absence of any external stress. The bulk chromatin association of 53BP1 shows dramatic dependence on PRIM1, which synthesizes the RNA primer of Okazaki fragments. The direct recruitment of 53BP1 to nascent DNA shows susceptibility to *in situ* ribonuclease A treatment. Conversely, depletion of FEN1, which results in the accumulation of uncleaved RNA primers, leads to an upregulation of 53BP1 levels at the replication forks, suggesting that RNA primers contribute to the recruitment of 53BP1 at the lagging DNA strand. 53BP1 depletion induces an accumulation of S phase poly(ADP-ribose), which constitutes a sensor of unligated Okazaki fragments. Collectively, our data indicate that 53BP1, distinct from its canonical mode of chromatin-binding, is anchored at the replication fork through its RNA-binding activity, highlighting the role of an RNA-protein interaction at DNA replication forks.

## INTRODUCTION

Recent observations highlight the importance of RNA-binding proteins (RBPs) in genome maintenance. RBPs, through their direct and generally specific binding to *cis*-acting elements in mRNAs, usually act as post-transcriptional regulators of gene expression ^1^. Importantly, in addition to their proven role in DNA damage-induced post-transcriptional regulation of DDR (DNA Damage Response) gene expression, RBPs, and the RNAs to which they bind, may play a more direct role in genome integrity ^2–4^. Several proteins involved in the DDR have been suggested to be RBPs. However, a potential RNA-binding activity of these proteins remains to be evaluated.

53BP1 (p53-binding protein 1) is a key protein that mediates the signaling of DNA double strand breaks (DSBs) in G1 and S/G2-phases of the cell cycle. In G1, 53BP1 acts as a molecular scaffold that recruits additional DSB-responsive proteins to damaged chromatin to limit DNA end-resection and promote repair by Non-Homologous End Joining (NHEJ) ^5^. The accumulation of 53BP1 at DSBs is RNA-dependent ^6–8^ and RNA immunoprecipitation (RIP) assays have shown that 53BP1 associates with RNAs ^7,8^. These observations however do not demonstrate a direct interaction between 53BP1 and RNA, even if a recent proteomic analysis has identified 53BP1 as a candidate protein interacting with RNA ^9^. In addition to its role in the DDR, 53BP1 plays an important function in DNA replication. Indeed 53BP1 has been shown to be rapidly recruited to stalled replication forks following a replicative stress induced by hydroxyurea (HU) treatment ^10^. Lack of 53BP1 decreases the cell survival and enhances chromosomal aberration after replication arrest ^11^. 53BP1 protects replication forks during the cellular response to replication stress in primary B cells ^12^ and the *S. cerevisiae* 53BP1 ortholog (RAD9) protects stalled replication forks from degradation in Mec1 (ATR) defective cells ^13^. The mechanism of 53BP1 recruitment to replication forks may not be completely similar to the one existing for 53BP1 recruitment to DSBs since depletion of RNF8, RNF168 or 53BP1, but not MDC1, displays a similar replication defect (*i*.*e*. delayed fork progression, and reversed fork accumulation) ^14^. It is therefore not clear which interaction(s) 53BP1 establishes to mediate its functions in DNA replication.

During initiation of DNA replication, several ribonucleotides are polymerized by the combined action of an RNA polymerase (DNA primase) and an error-prone DNA polymerase (POLα). DNA polymerases are unable to initiate DNA synthesis *de novo* and require an initiating step to generate an RNA primer. This “priming” role is fulfilled by specialized DNA-dependent RNA polymerases, called DNA primases containing a small catalytic subunit (PRIM1) and a large accessory subunit (PRIM2), capable of synthesizing short RNA chains (7-12 ribonucleotides). Primases are also essential throughout the process of lagging strand replication, where they initiate synthesis of the discontinuously synthesized Okazaki fragments ^15^. On the lagging strand, the DNA polymerase δ (POLδ) synthesizes approximately 100 deoxyribonucleotides downstream of this RNA-DNA primers, to form the Okazaki fragments ^16^. When POLδ displaces the downstream fragment, it creates a short 5’ flap single-stranded nucleic acid that is cleaved by the FEN1 endonuclease to allow subsequent ligation of the two Okazaki fragments by DNA ligase I (LIG1). In some instances, a long flap is created that is covered with RPA thereby blocking the access to FEN1. RPA can then recruit the DNA nuclease/helicase DNA2, which cleaves the long flap into a short flap that can be subsequently cleaved by FEN1 ^17–20^. Whether RBPs, reported to be present at replication forks ^21^, directly bind RNA primers of Okazaki fragments during unperturbed DNA replication is unknown.

Using CLIP (UV-C induced CrossLinking ImmunoPrecipitation ^22^) and 2C (Complex Capture ^23^) experiments in living cells, we provide evidence that 53BP1 directly interacts with RNA. By analysing the nature of the nucleic acids bound to 53BP1, we found entities of about 20-200 nucleotides, composed of ribonucleotides at the 5’ end followed by deoxyribonucleotides at the 3’ end, that constitute Okazaki fragments. Consistent with the hypothesis that 53BP1 directly interacts with Okazaki fragments, we found that RNAse treatment decreases 53BP1 recruitment to the fork (as determined by *in situ* analysis of protein interactions at DNA replication forks (SIRF ^24^). In addition, while depletion of PRIM1 leads to reduced 53BP1 assembly, loss of FEN1 enhances the 53BP1 association with nascent DNA. Finally, depletion of 53BP1 leads to reduced RPA and DNA2 loading at nascent DNA. Altogether these results indicate that the interaction between 53BP1 and Okazaki fragments is a key determinant of their processing, and more generally of the functions of 53BP1 in DNA replication.

## RESULTS

### 53BP1 is an RNA-Binding Protein

53BP1 contains a putative intrinsically disordered GAR (Glycin-Arginin Rich) RNA-binding domain ^25^ (Supplemental Figure 1A). To investigate a potential RNA-binding activity, we first used an electrophoretic mobility shift assay (EMSA). The migration of the structured *SC1* RNA (Supplemental Figure 1B) (known to interact with the GAR domain of the FMRP protein; Supplemental Figure 1C) was shifted by a recombinant GAR-Tudor 53BP1 protein (Supplemental Figure 1D and E). Surface Plasmon Resonance (SPR) experiments indicated that the GAR-Tudor fragment interacts with the *SC1* RNA with an affinity of about 8 nM, while a GAR-Tudor fragment mutated in the GAR domain (3R to 3K) completely lost its affinity with the *SC1* RNA (Supplemental Figure 1F). The ability of 53BP1 to directly bind to RNAs *in vivo* was analyzed following irradiation of living cells with high doses of UV (254 nm) in a CLIP (CrossLinking ImmunoPrecipitation) experiment ^22^. UV crosslinking requires direct contact (“zero” distance) between protein and RNA and does not promote protein-protein crosslinking ^26,27^. We irradiated human HEK293T living cells with UV-C and immediately prepared cell lysates. Following 53BP1 immunoprecipitation and stringent washes, the immunoprecipitate (IP) was radiolabeled with T4 polynucleotide kinase (PNK) that incorporates ^32^P at the 5’-end of nucleic acids (Figure 1A and B). In UV-exposed cells, we observed a sharp radioactive band at the size of 53BP1 (Figure 1B), indicating that 53BP1 directly interacts with RNAs. An siRNA pool directed against 53BP1 was used to ascertain that the band migrating at the size of 53BP1 corresponds to 53BP1 and that the lower band (labelled *) corresponds to a non-specific signal. We obtained very similar results in other cell lines, including Hela cervical cancer cells, A2058 melanoma cells, RPE1 non-transformed human epithelial cells and U2OS osteosarcoma cells (Supplemental Figure 2A). To ascertain that the above CLIP results were not linked to an artefact related to the use of the 53BP1 antibody, we expressed GFP-53BP1 fusion proteins in HEK293T cells and performed the CLIP experiment with a GFP antibody, instead of the 53BP1 antibody. A radioactive band was observed at the predicted size of the GFP-53BP1 protein or of the GFP-53BP1-Cter protein containing the carboxy-terminal half of 53BP1 that is still able to be recruited to ionizing radiation-induced damaged DNA (Supplemental Figure 2B and C). Altogether, these data show that an RNA-binding activity can be found in the carboxy-terminal half of 53BP1 containing the GAR-Tudor domain.

**Figure 1.**
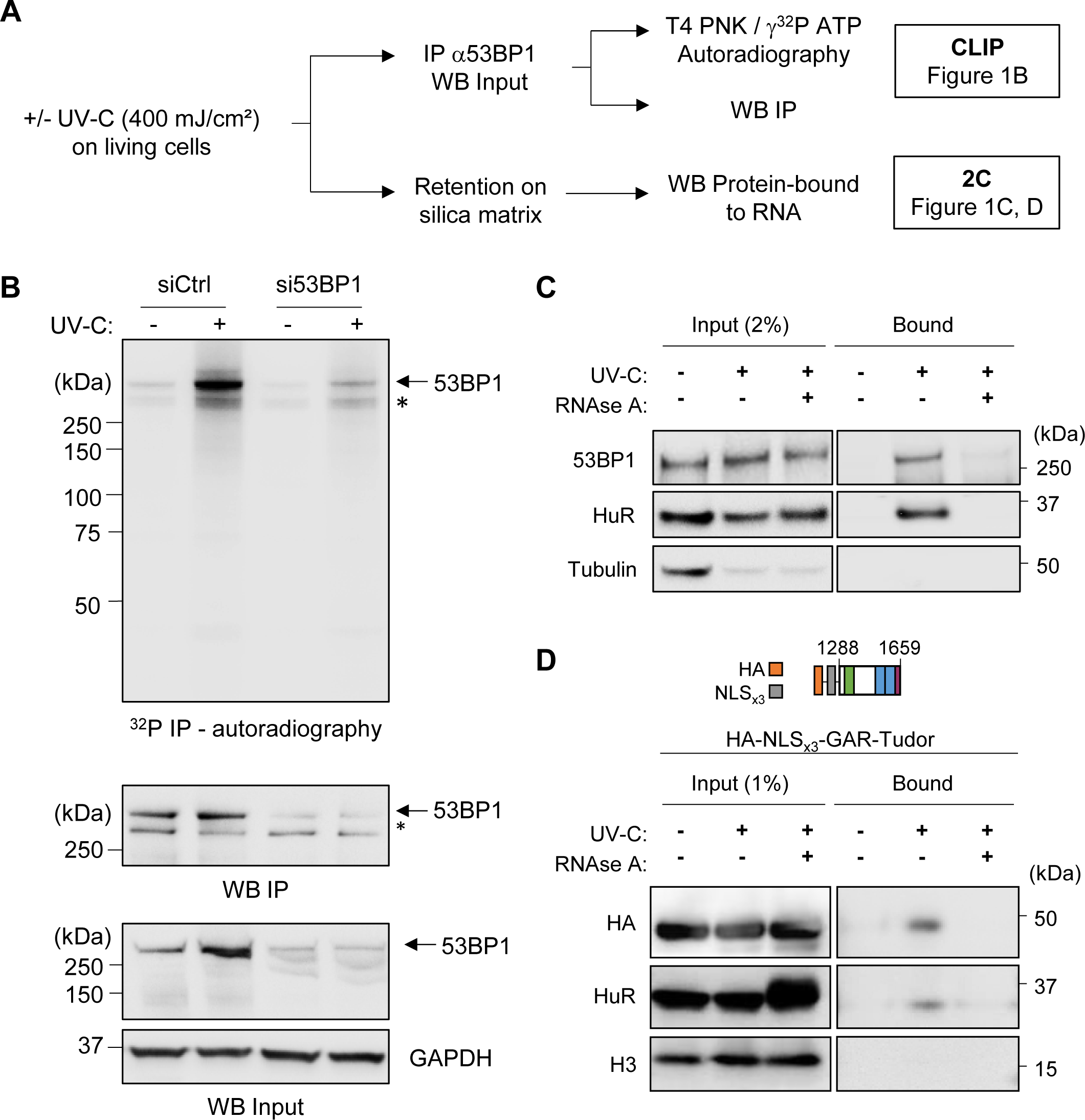
53BP1 directly interacts with RNA. (A) Schematic representation of the CrossLinking ImmunoPrecipitation (CLIP) and Complex Capture (2C) procedures. WB: western blot; IP: immunoprecipitation. PNK: polynucleotide kinase. (B) Upper panel: Autoradiography of 53BP1-nucleic acids complexes in UV-C treated (+) or untreated (-) HEK293T cells depleted of 53BP1 (si53BP1) or left untreated (siCtrl). Lower panel: IP of 53BP1 and WB of 53BP1 and GAPDH (used as a normalization control); same conditions as in upper panel. The asterisk indicates a non-specific band (*), the arrow indicates the position of 53BP1-nucleic acid complexes (n=3). (C) WB on 2C experiments performed in A2058 cells (n=3). (D) Western blot on 2C experiments performed in GAR-Tudor transfected HEK293T (n=3).

Even if UV irradiation is mainly used to monitor RNA-protein interactions ^22^, it has also been used to establish crosslinks between proteins and single-stranded DNA (ssDNA) (*e*.*g*. Steen et al., 2001). We therefore used an orthogonal approach (complex capture -2C) that uses the property of silica matrices to strongly and specifically retain nucleic acids and crosslinked nucleic acid-protein complexes based on charges ^23^ (Figure 1A, C and D). 53BP1 was retained on the silica matrix in a UV- and RNA-dependent manner (Figure 1C). The well-characterized RBP HuR and the tubulin were used as positive and negative controls, respectively. Consistent with the EMSA data, the GAR-Tudor region is sufficient to interact with RNAs in living cells (Figure 1D). Altogether, these data show that 53BP1 directly interacts with RNA *in vitro* and *in vivo*.

### 53BP1 binds RNA-DNA chimeras

To analyse the nature of the nucleic acids bound to 53BP1, we extracted nucleic acids from the ^32^P-labelled crosslinked nucleic acid-GFP-53BP1 complexes, loaded them on a denaturing TBE-Urea gel and revealed the radioactive signal by autoradiography. We found that nucleic acids bound to 53BP1 migrate as a smear of about 20-150 nucleotides (Figure 2A). The smear collapsed below 25 nucleotides following treatment with ribonuclease A (RNAse A; which in our experimental conditions cleaves both single-stranded and double-stranded RNA, but not DNA or RNA engaged in an RNA-DNA duplex; Supplemental Figure 3) showing that these nucleic acids possess ribonucleotides at their 5’ ends. Strikingly, the size of the smear was also shortened following treatment with deoxyribonuclease I (DNAse I) (which specifically cleaves DNA, but not RNA; Supplemental Figure 3), indicating that the nucleic acids interacting with 53BP1 are also composed of deoxyribonucleotides. The remaining DNAse I-resistant shorter smear was composed of RNA at its 5’-end since it disappeared following treatment with both RNAse A and DNAse I (Figure 2A). Similar results were found with the endogenous 53BP1 (Supplemental Figure 4A). DNAse I had no effect on nucleic acids bound to the RNA-binding protein RBMX (Figure 2B), ruling out the possibility that DNAse I was contaminated with RNAses. We concluded that the cellular nucleic acids interacting with 53BP1 are RNA-DNA chimeras composed of 5’-end RNAs of different sizes (less than 25 nts) followed by deoxyribonucleotides (up to 100 nts) at their 3’ ends. Of note, the GAR-Tudor fragment that is sufficient to bind RNA (Figure 1D and Supplemental Figure 1) cannot bind RNA-DNA chimeras since DNAse I treatment didn’t lead (in contrast to RNAse treatment) to a reduction in the size of the smear (Supplemental Figure 4B) that is therefore only composed of ribonucleotides. This indicates that the binding of 53BP1 to Okazaki fragments through the GAR domain requires additional 53BP1 protein domains.

**Figure 2.**
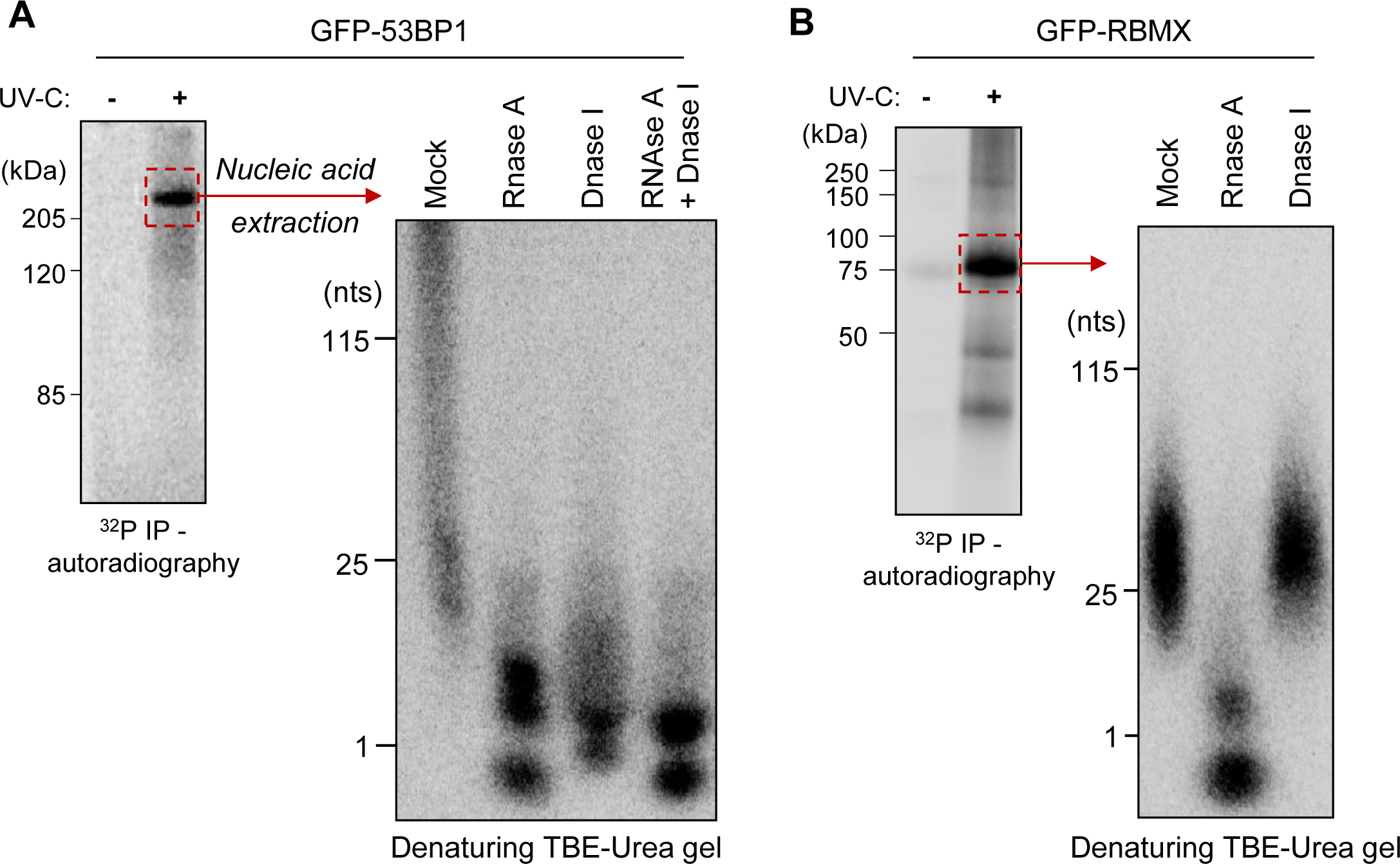
53BP1 binds an RNA-DNA chimera. CLIP (left panel) and nucleic acid extraction (right panel) from GFP-53BP1 (A) or GFP-RBMX (B) transfected HEK293T (n=3).

### 53BP1 interacts with Okazaki fragments

Based on these findings, we hypothesized that Okazaki fragments (the only known RNA-DNA chimeras in human cells) might contribute to the recruitment of 53BP1 to DNA replication forks. To test this hypothesis, we depleted the human PRIM1 subunit that synthesizes RNA primers of replication and employed quantitative image-based cytometry (QIBC) ^29^ to quantify the chromatin abundance of 53BP1 at the single-cell level. The depletion of PRIM1 with two independent siRNAs led to a decrease of 53BP1 association with chromatin (Figure 3A and Supplemental Figure 5A) in the different phases of the cell cycle (Supplemental Figure 5B) without altering the cell cycle profile (Supplemental Figure 5C and Supplemental Figure 5D). Consistent with previous results ^30^, PRIM1 loss led to DNA replication stress, as seen from the pronounced induction of ssDNA and chromatin loading of RPA2 during the S-phase (Supplemental Figure 5E and Supplemental Figure 5F). Interestingly, while PRIM1 depletion decreased the global chromatin association of 53BP1 (Supplemental Figure 6A), it had a modest effect of disturbing spontaneously arising 53BP1 foci assembled on chromatin due to dedicated DDR signaling (Supplemental Figure 6B). Together, these results suggest that in addition to the well-established histone modification-based chromatin loading of 53BP1 at DSBs ^5^, RNA primers synthesized by PRIM1 could facilitate the recruitment of 53BP1 on chromatin during DNA replication.

**Figure 3.**
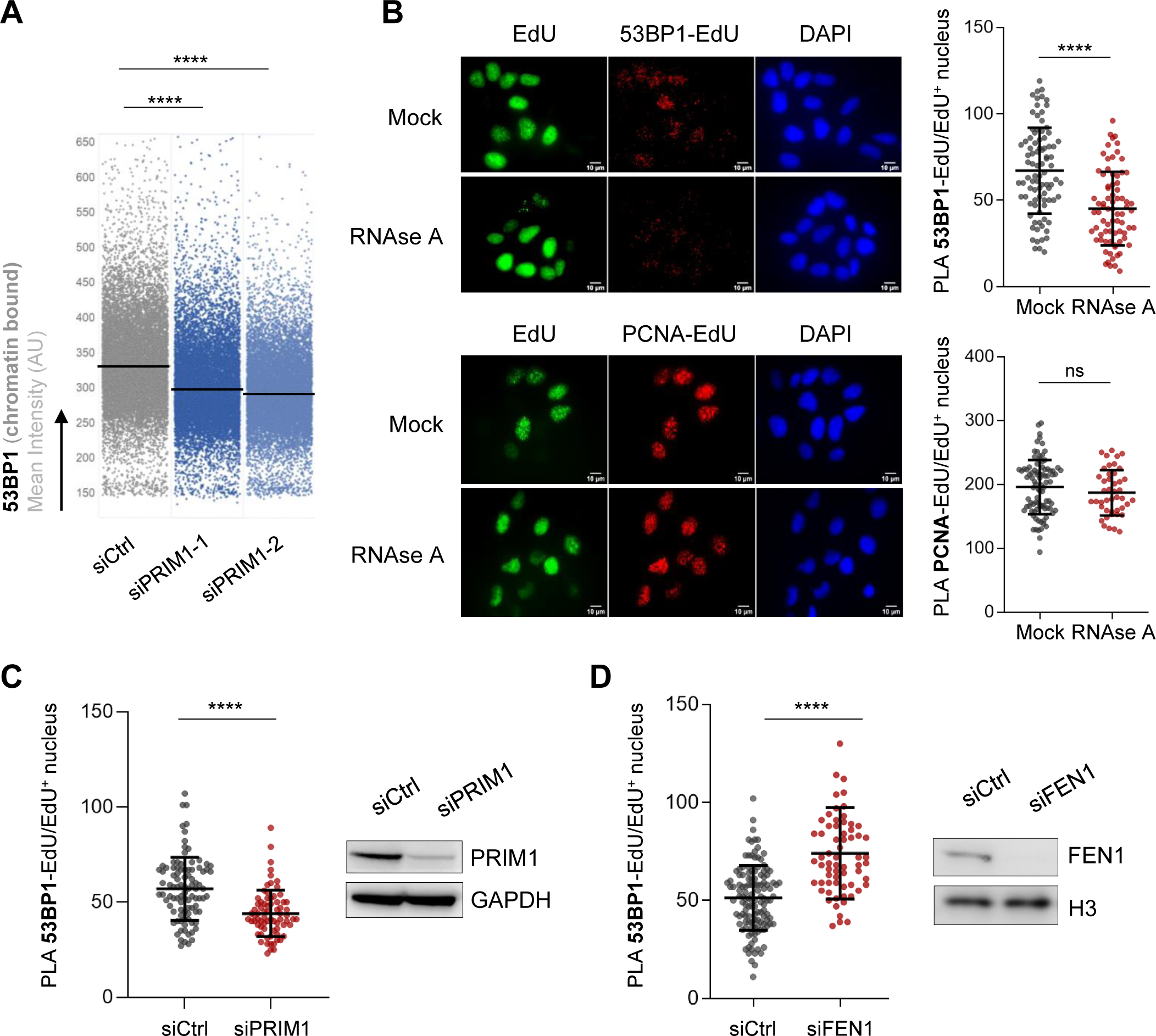
53BP1 interacts with the RNA primer of Okazaki fragments. (A) QIBC of chromatin loaded 53BP1 mean intensity. U2OS cells were treated with indicated siRNAs for 48 hours (also see Figure. S6). The horizontal lines are median values. P values were determined by one-way ANOVA with Tukey’s test, **** p<0.0001 (n > 10,000 cells from 3 biological repeats). (B) Representative images and/or analysis of 53BP1-EdU and PCNA-EdU SIRF in U2OS cells permeabilized and treated with RNAse A (RNAse A) or left untreated (Mock); PLA signal (red), Alexa Fluor 488–EdU staining (green), and DAPI staining (blue). The significance for 53BP1-EdU and PCNA-EdU PLA values (shown as a scatter plot) was derived from the Mann-Whitney statistical test. Bars represent the mean ± s.d. **** p<0.0001 (n=3). (C-D) Representative scatter plot of U2OS cells depleted of PRIM1 (siPRIM1) (C), FEN1 (siFEN1) (D) or left untreated (siCtrl) for 48 hours. The significance was derived from the Mann-Whitney statistical test. Bars represent the mean ± s.d. **** p<0.0001. (n=3).

While the QIBC approach allows the determination of the chromatin abundance of proteins, it does not allow the determination of the abundance of proteins at replicating DNA. We therefore used an *in situ* analysis of protein association at DNA replication forks (SIRF) procedure that allows for the quantitative assessment of the proximity of proteins with ongoing, stalled, and previously active replication forks using a Proximity Ligation Assay (PLA) approach ^24^. As expected, 53BP1 could be found associated with nascent DNA (Supplemental Figure 7). Treatment of cells with RNAse A (Figure 3B) led to a decreased proximity between 53BP1, but not PCNA (a component of the replisome), and nascent DNA, without changes in EdU incorporation (Supplemental Figure 8A and Supplemental Figure 8B). The same result was observed upon depletion of PRIM1 (Figure 3C and Supplemental Figure 8C). In contrast, depletion of FEN1, that leads to an accumulation of unligated Okazaki fragments ^20^, led to an increased proximity between 53BP1 and nascent DNA (Figure 3D and Supplemental Figure 8D). Taken together, these data indicate that perturbation of the synthesis and maturation of Okazaki fragments through depletion of PRIM1/FEN1 or RNA degradation leads to reduced proximity between 53BP1 and nascent DNA.

### Depletion of 53BP1 perturbs the lagging strand processing

Okazaki fragment maturation during lagging strand replication involves several layers of nuclease-driven pathways for proper RNA–DNA primer removal. Two pathways exist, the short flap pathway mediated by FEN1 and the long flap pathway mediated by DNA2/RPA. Based on our results that 53BP1 associates with Okazaki fragments during active DNA synthesis, we next sought to determine whether 53BP1 binding to RNA–DNA primers plays a hitherto unknown role in supporting Okazaki fragment maturation. We found that 53BP1 depletion leads to an increased accumulation of mono- and poly-(ADP-ribose) (MAR, PAR) in chromatin (Figure 4A and Supplemental Figure 9) that is more pronounced in S and G2 (Supplemental Figure 9B) and which constitutes a sensor of unligated Okazaki fragments ^31^. Short incubation with PARG inhibitor to preserve nascent PAR further enhanced the levels of ADP ribosylation in 53BP1 depleted cells (Figure 4A). We also found that depleting 53BP1 leads to reduced RPA and DNA2 loading at chromatin (Figure 4B, C, D and Supplemental Figure 10A-D), as well as accumulation of single-stranded DNA (Supplemental Figure 10E). Collectively, these results indicate that 53BP1 binds to long flaps of Okazaki fragment, fosters proper RNA–DNA primer removal and prevents genotoxic DNA lesions that might arise from the lagging DNA strands in the absence of external stress inducer.

**Figure 4.**
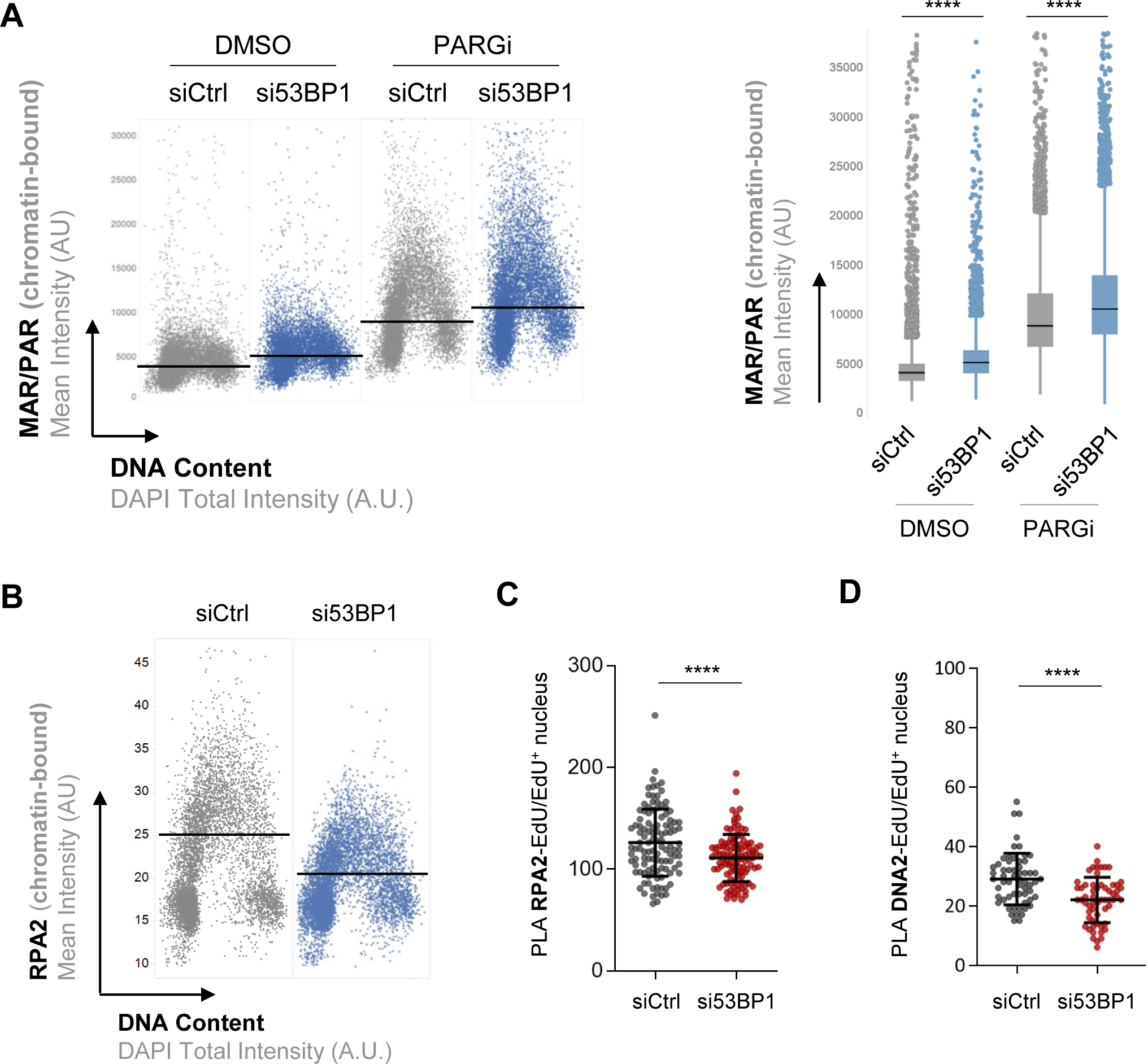
Depletion of 53BP1 perturbs the lagging strand processing. (A) QIBC of chromatin loaded protein mono ADP-ribosylation (MARylation) or poly ADP-ribosylation (PARylation). U2OS cells were treated with either control or 53BP1 siRNAs for 72 hours, and incubated with DMSO or PARGi (10 µM) for the last 30 minutes before pre-extraction and cell fixation. Nuclear DNA was counterstained by DAPI. The horizontal lines are medians. Right: quantification of MAR/PAR signals from the experiment in Left. In box plots, centre lines are medians, the boxes indicate the 25th and 75th centiles, the whiskers indicate Tukey values. P values were determined by one-way ANOVA with Tukey’s test, **** p<0.0001 (n > 10,000 cells from 2 biological repeats). (B) QIBC of chromatin loaded RPA2 mean intensity. U2OS cells were treated with either control or 53BP1 siRNAs for 72 hours. The horizontal lines are medians (n > 5,000 cells from 2 biological repeats). (C-D) Representative scatter plot of SIRF experiments of RPA (C) or DNA2 (D) in U2OS cells depleted of 53BP1 (si53BP1) or left untreated (siCtrl) for 48 hours. The significance was derived from the Mann-Whitney statistical test. Bars represent the mean ± s.d. **** p<0.0001. (n=3).

## DISCUSSION

Here we show that 53BP1 directly interacts with RNA *in vitro* and *in vivo* in human cells, revealing a new activity of 53BP1 as a *bona fide* RNA-binding protein (Figure 1). Strikingly, the nucleic acids interacting with 53BP1 are RNA-DNA chimeric structures composed of 5’-end ribonucleotides followed by deoxyribonucleotides at their 3’ ends (Figure 2) that constitute Okazaki fragments (Figure 3). 53BP1, distinct from its canonical mode of chromatin-binding, is anchored at the replication fork via its RNA-binding activity, thereby constituting the first ever-described binder of Okazaki fragments during unperturbed DNA replication. Of note it has been recently established that, during replication stress in fission yeast, the NHEJ factor Ku, also known as an RBP ^4^, binds to RNA:DNA hybrids originating from Okazaki fragments and establishes a barrier to replication fork degradation ^32^. Our findings show that while the GAR-Tudor domain of 53BP1 is sufficient to bind RNA, it is not able to bind RNA-DNA chimeras in living cells. This indicates that binding of 53BP1 to RNA-DNA chimeras requires an additional RNA/DNA binding domain, an additional 53BP1 domain that recruits the protein in the vicinity of Okazaki fragments or that the GAR-Tudor domain alone adopts a conformation that affects its binding to RNA-DNA chimeras. Ideally, structural biology approaches will facilitate the defining of the minimal 53BP1 domain able to interact with RNA-DNA chimeras and an associated mutant. Moreover, even if our data indicates that 53BP1 interacts with Okazaki fragments on the lagging strand, the binding of 53BP1 to any RNA-DNA primers including the one present on the leading strand cannot be completely ruled out.

Similar to its role in DNA double-strand break repair pathway choice ^5^, 53BP1 may contribute to the choice between the short and long flap pathways during Okazaki fragment maturation. In yeast it has been show that a delay in cleavage by FEN1 can induce a long flap ^18^. Our data on the association of RPA and DNA2 with nascent DNA upon 53BP1 depletion (Figure 4) indicate that the interaction between 53BP1 and the long flap may contribute to prevent the action of FEN1, thus supporting the long flap pathway and further recruitment of RPA/DNA2. Maturation through the long flap pathway was shown to limit mutation rates, especially in transcribed regions, because of the replacement of the DNA primer synthesized by the error-prone POLα with a DNA strand synthesized by the error-free POLδ ^33,34^. In this scenario, 53BP1 may contribute to maintain genome integrity during DNA replication.

This new role of 53BP1 in the vicinity of replication fork might be of prime importance in human health. Indeed, defects in the maturation of Okazaki fragments lead to increased genetic instability, which causes certain cancers ^35–37^. Moreover, several studies have indicated that *TP53BP1* mutation might be associated with cancer risk ^38,39^. Hence, the evaluation of 53BP1 mutations should be extended, beyond their effects on DSBs signaling, to replication defects. In addition, it will be interesting to analyse the role of the RNA-binding activity of 53BP1 in the context of the sensitivity to poly(ADP-ribose) polymerase inhibitors (PARPi) of tumor cells, that are deficient in hereditary breast cancer genes (BRCA1 and BRCA2). PARPi, that has anticancer activity in BRCA-deficient cancers, perturbs the processing of Okazaki fragments, a process that is modulated by 53BP1 loss ^40–42^. The Okazaki fragment binding activity of 53BP1 may specifically be involved in the sensitivity of BRCA1-deficient cells to PARPi and linked to DNA replication gaps.

## MATERIAL AND METHODS

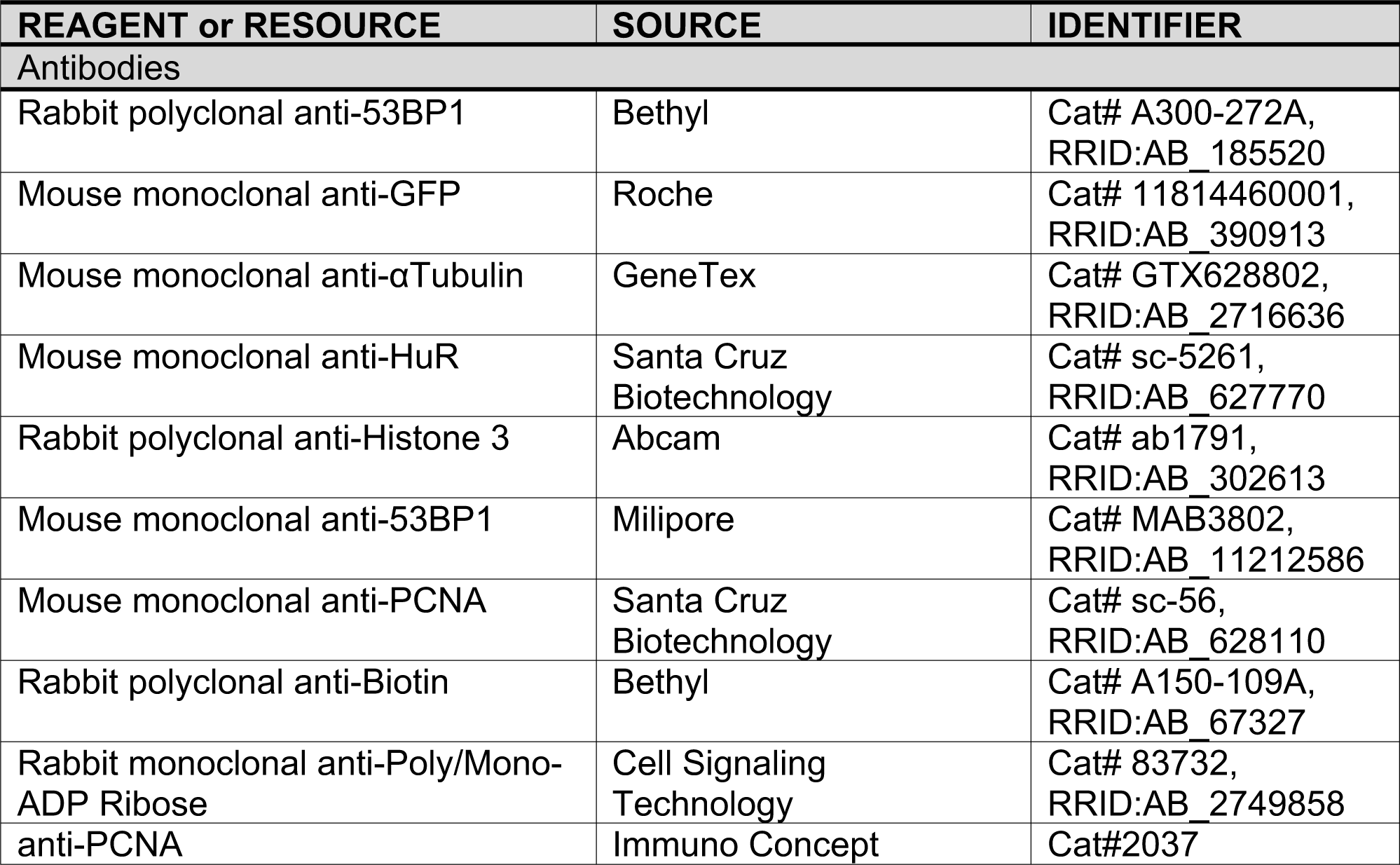
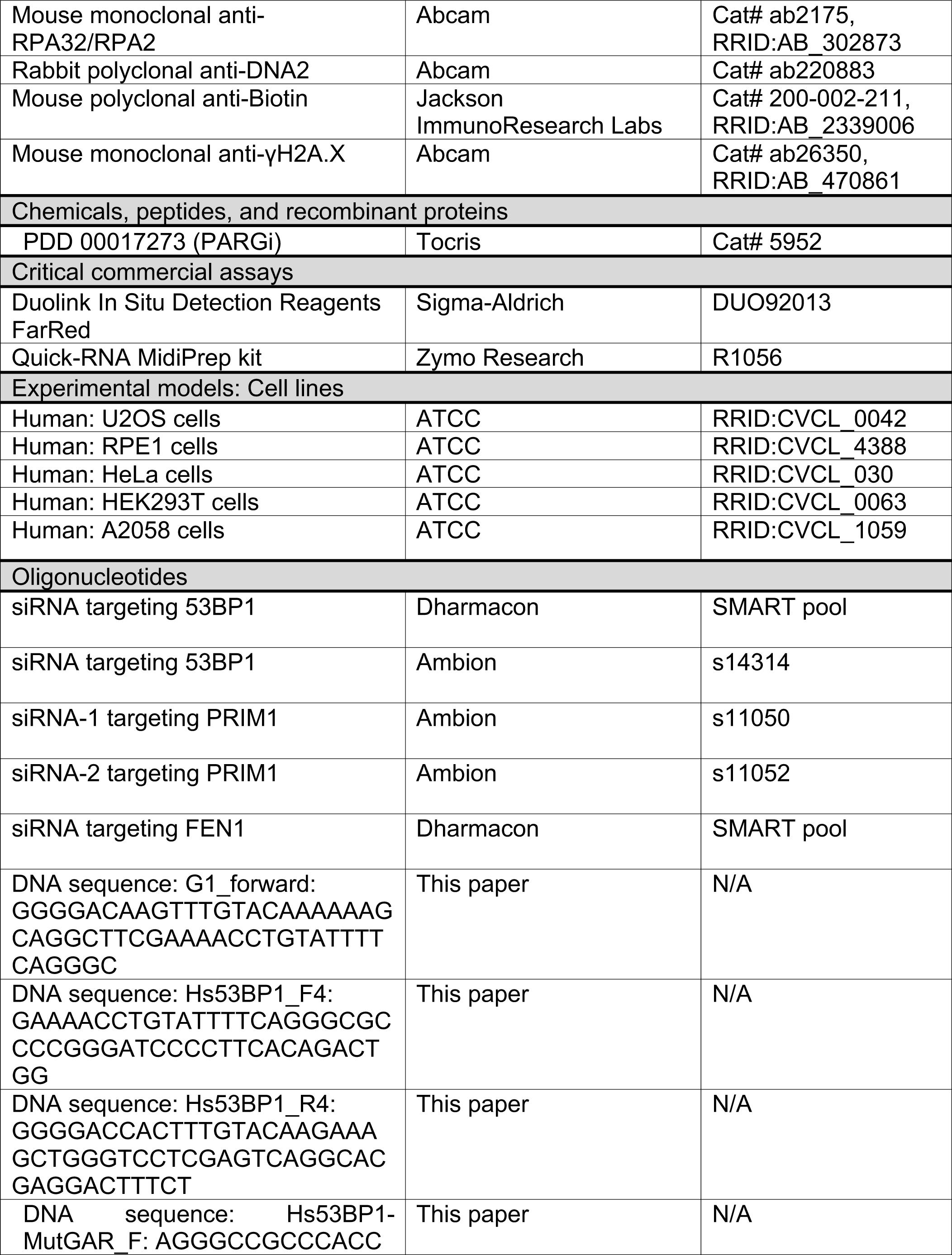
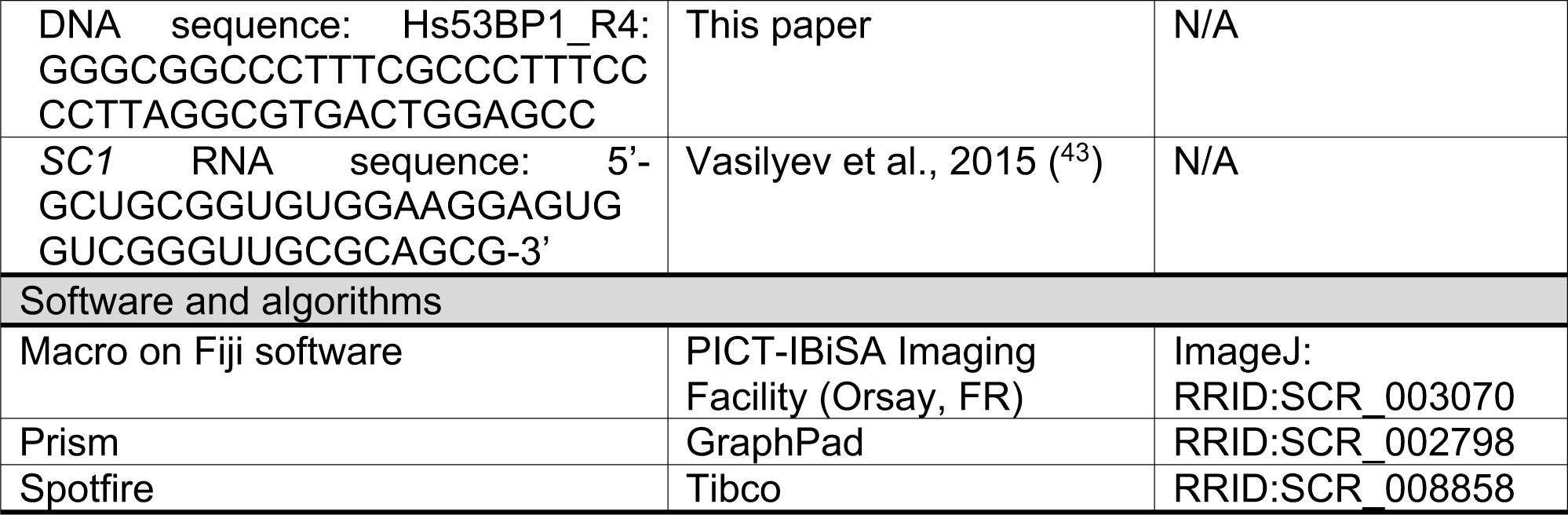

### Cell culture

Cells were grown in 5% CO_2_ humidified incubator at 37°C. U2OS, RPE1, HeLa and HEK293T cells were grown in DMEM supplemented with 10% FBS and 2 mM L-glutamine. A2058 cells were grown in MEM supplemented with 10% FBS and 2 mM L-glutamine.

### siRNA and Plasmid transfections

siRNA and plasmid transfections were performed, respectively, using Lipofectamine RNAiMAX (Life Technologies, #13778150) or JetPEI (Polyplus Transfection, #101000053) according to manufacturer’s instructions.

### CrossLinking and immunoPrecipitation (CLIP)

Cells were irradiated with UV-C light at 254nm, lysed in RIPA buffer (50 mM Tris-HCl pH 7.4, 100 mM NaCl, 1% Nonidet P-40, 0.1% SDS, 0.5% Sodium deoxycholate), supplemented with RNAse OUT (1µL/1mL, Invitrogen, #10777019) and protease inhibitor cocktail, and immunoprecipitated with 50 µL of Dynabead Protein G (Thermofisher, #10009D) using 5 µg of rabbit anti-53BP1 or 5 µg of mouse anti-GFP antibodies for each condition. Immunocomplexes were washed twice with RIPA-S buffer (50 mM Tris-HCl pH 7.4, 1 M NaCl, 1 mM EDTA, 1% Nonidet P-40, 0.1% SDS, 0.5% Sodium deoxycholate) and once with PNK buffer (20 mM Tris-HCl pH 7.4, 10 mM MgCl_2_, 0.2% Tween 20). Then, crosslinked nucleic acids were radiolabelled using ATP, [γ-32P] (PerkinElmer, #BLU502A250UC) and T4 polynucleotide kinase (PNK, Thermofisher, #EK0032), washed twice with RIPA-S buffer and once with PNK buffer.

Protein-RNA complexes are eluted and separated by SDS-PAGE. Then the gel was dried and complexes were visualized by exposure to a phosphoimager screen. In the case where nucleic acids were extracted, RNA-proteins complexes were transferred onto nitrocellulose membrane 0.45 µm and complexes were visualized by exposure to a phosphoimager screen. The nitrocellulose membrane was cut out at the suspected molecular weight of the radiolabelled protein of interest. Nitrocellulose pieces were treated with proteinase K (1 mg/mL, ThermoScientific, #EO0491) in 200 µL of PK buffer (100 mM Tris-HCl pH 7.4, 50 mM NaCl, 10 mM EDTA), once without urea and then once with urea 7 M. Nucleic acids were purified with phenol/chloroform/isoamylic alcohol 25:24:1 (Sigma) and then precipitated overnight by adding Glycoblue (Thermofischer, #AM9515), sodium acetate and 100% ethanol. After centrifugation and washes (80% ethanol), nucleic acids pellet was resuspended in water and divided into three conditions. One condition without treatment, one with RNAse A treatment (1 µL at 10 mg/mL, ThermoScientific, #EN0531) and one with DNAse treatment (1µL at 2U/µL, Invitrogen, #AM2238). Nucleic acids were separated using denaturing polyacrylamide gel (TBE-Urea gel, 6%). The gel was dried and nucleic acids were visualized by exposure to a phosphoimager screen.

### Complex capture (2C)

Cells were irradiated with UV-C light at 254 nm and lysed in HMGN150 buffer (20 mM Hepes pH 7,5; 150 mM NaCl; 2 mM MgCl_2_; 0.5% Nonidet P-40; 10% Glycerol). The lysate is then treated or not with RNAse A (15µL for 1 mg of proteins). A fraction (2%) of the input is kept to load on a SDS-PAGE to control the amount of proteins. The quick-RNA Midiprep kit is used for RNA extraction for the 2C Method. The RNA concentration of the eluate is measured using NanoDrop (ThermoScientific). 50µg RNA is treated with RNAse I (500U, Invitrogen, #AM2295) for 40 min at 30°C. A mix of LDS 4X (Thermofischer, #NP0007) + Reducing reagent 10X (Thermofischer, #NP0009) are added to the samples and a western blot analysis is performed.

### Quantitative image-based cytometry (QIBC)

QIBC was performed as previously described ^44^. Briefly, images were acquired with a ScanR inverted microscope high-content screening station (Olympus, IX81) equipped with wide-field optics, air objective, fast excitation and emission filter-wheel devices for DAPI, FITC, Cy3, and Cy5 wavelengths, an MT20 illumination system, and a digital monochrome Hamamatsu ORCA-Flash 4.0LT CCD camera. Images were acquired in an automated fashion with the ScanR acquisition software (Olympus, 3.2.1). 81-100 images were acquired containing at least 5,000 cells per condition. Acquisition times for the different channels were adjusted for non-saturated conditions in a 12-bit dynamic range, and identical settings were applied to all the samples within one experiment. Images were processed and analysed with ScanR analysis software. First, a dynamic background correction was applied to all images. The DAPI (Sigma-Aldrich) signal was then used to generate an intensity-threshold-based mask to identify individual nuclei as the main objects. This mask was then applied to analyse pixel intensities in different channels for each individual nucleus. For analysis of 53BP1 foci, additional masks were generated by segmentation of individual 53BP1 spots with spot-detector modules included in the software. Each focus was defined as a subobject, and this mask was used to quantify pixel intensities in foci. After this segmentation of objects and subobjects, the desired parameters for the different nuclei or foci were quantified with single parameters (mean and total intensities, area, foci count, and foci intensities).

These values were then exported and analysed with TIBCO Spotfire Software, version 11.1, to quantify absolute, median, and average values in cell populations and to generate all color-coded scatter plots. Within one experiment, similar cell numbers were compared for the different conditions (at least 4,000–5,000 cells), and for visualization, low x-axis jittering was applied (random displacement of objects along the x-axis) to make overlapping markers visible.

### *In situ* protein interaction with nascent DNA replication forks (SIRF)

U2OS were seeded on Millicell EZ SLIDE 8-well glass (Millipore, #PEZGS0816) 48h before experiment and then incubated with 25 µM EdU for 10 minutes. After treatment, cells were washed with cold PBS, pre-extracted with cold CSK (10 mM PIPES, 100 mM NaCl, 300 mM sucrose, 3 mM MgCl_2_, protease inhibitor cocktail), permeabilized with cold CSK-T buffer (CSK buffer supplemented with 2% Triton X-100) during 5 minutes at 4°C, treated or not with 1mg/mL of RNAse A for 5 min at 37°C and then fixed with 4% PFA in PBS for 20 minutes at room temperature. After fixation, slides were placed in a humid chamber, and incubated with a click-it reaction (10 µM biotin azide, 100 mM sodium ascorbate, 2 mM CuSO_4_, 1 µM Alexa Fluor 488 Azide, in PBS in this order) at room temperature for 1 hour. After the click reaction, slides were washed with PBS and blocked with blocking buffer (10% goat serum and 0.1% Triton X-100 in PBS) for 1 hour at room temperature. Primary antibodies (anti-53BP1, anti-biotin, anti-PCNA) were diluted in blocking buffer and incubated at 4°C overnight. Then a PLA procedure is performed using Duolink In situ Detection Reagents FarRed. Slides were imaged using Leica 3D upright deconvolution microscope with a CoolSNAP HQ camera, at the PICT-IBiSA Imaging Facility in Orsay, and analysed using a semi-automatic macro on Fiji software.

### Cloning

GFP-RBMX and GFP-53BP1 plasmids correspond to RBMX or 53BP1 sequence inserted in pcDNA 6.2 (Vivid Colors(tm) pcDNA(tm)6.2/N-EmGFP-GW/TOPO(tm) Mammalian Expression Vector, Thermofisher). The GFP-53BP1-Cter corresponds to the 1235 to 1972 encoding sequence of 53BP1.

The Hs53BP1-1288-1659 (HA-NLSx3-GAR-Tudor) encoding sequence was cloned by site-directed recombination using the Gateway technology (Life Technologies). The ORF was amplified by PCR using specific primers: G1_forward; Hs53BP1_F4; Hs53BP1_R4.

The resulting PCR products were recombined during BP reaction with pDONO207 to generate Entry clones used for further LR reactions with pGGWA ^45^.

The triple mutation (Arg1396Lys, Arg1398Lys, Arg1401Lys) was generated by “rolling-circle” PCR using specific primers as previously described ^46^: Hs53BP1-MutGAR_F and Hs53BP1-MutGAR_R.

### 53BP1 fragments purification

GST-Hs53BP1-1288-1659 fragments (GAR-Tudor or GAR(3K)-Tudor) purifications were performed as described ^47^. Fractions eluted from Superdex 75 10/300 GL (Cytiva) were directly subjected to TEV cleavage at the ratio of 1 TEV for 10 GST-Hs53BP1 fragment (w/w). After 2 hours incubation at 20°C, the mixture was incubated with Glutathione Sepharose™ 4B (Cytiva) and Ni-NTA Superflow (Qiagen). The Hs53BP1 fragment without its GST tag was recovered in the flow through and concentrated using Vivaspin concentrator (Sartorius).

### Electrophoretic Mobility Shift Assay (EMSA)

Appropriate concentrations of each protein and trace amount (25 nM) of ATTO700-labeled oligonucleotides were incubated with binding buffer (10 mM Tris-HCl, 1 mM EDTA, 100 mM KCl, 0.1mM DTT, 5% glycerol, 0.01 mg/mL BSA, pH 7.5) in a 20 µL final reaction volume at room temperature for 45 min. The incubated samples were loaded on an 8% polyacrylamide (37.5:1 acrylamide/bis-acrylamide) non-denaturing gel. Gels were imaged using an Odyssay scanner.

### Surface Plasmon Resonance (SPR)

The interaction between nucleic acids (ligands) and the proteins (analytes) were measured by the surface plasmon resonance (SPR) technique using a ProteOn XPR36TM (Bio-Rad Laboratories). NLC Sensor chips precoated with NeutrAvidin (Bio-Rad Laboratories) were conditioned with 30-μl injections of 50 mM NaOH and 1.0 M NaCl at a flow rate of 30 μl/min in both ligand and analyte flow directions. Immobilization was performed in the running buffer (10 mM Tris-HCl, 200 mM KCl at pH 7.5, Tris-HCl, 0.01% v/v Tween). The temperature of the chip surface was maintained at 25 °C. The all biotinylated oligonucleotides were immobilized one by one onto a single channel of the six channel NLC sensor chip. Each oligonucleotide construct was immobilized onto an individual ligand channel by flowing the constructs at a concentration of 10 to 40 nM for 60 to 240 sec injection. Binding to the chip surface resulted in approximately 60-100 response units (RU). The chip was rotated to the analyte flow direction and stabilized by flowing 60 μL analyte EMSA buffer three or more times, as required. Analyte (purified protein or buffer) was applied at various concentrations through the analyte channels at 100 μL/min for an association phase of 150 sec, followed by a dissociation phase (running buffer only) of 600 sec. The surface was regenerated with 18 μL of 0.5% SDS, at a flow rate of 30 μL/min followed by flowing 60 μL running buffer in the analyte direction at 100 μL/min. The resultant sensorgrams for each analyte interaction were analysed with ProteOn software (Bio-Rad Laboratories). Kinetic parameters for first the dissociation constant (kd) and then the association constant (ka) were estimated by fitting the kinetic data according to the 1 : 1 Langmuir model and values of KD were then calculated (using the ratio kd/ka).

### DNA damage induction before IF

Micro-irradiation: the confocal microscope TCS SP5 Leica allowed DNA damage induction in cultured living U2OS cells using laser at 405 nm.

### QIBC of native BrdU

To detect ssDNA in parental strands, the cells were labeled with 10 mM BrdU for 24 h and then released into a fresh growth medium for 1h prior to fixing cells. Cells were fixed, permeabilized, and processed for immunofluorescence staining as described in the IF and QIBC protocol. To detect BrdU in the native state, fixed cells were incubated with Rat monoclonal anti-BrdU antibody (BU1/75, ab6326; Abcam) for 90 min followed by Alexa-594 goat anti-Rat (A-11007; Thermo Fisher Scientific) secondary antibody at room temperature. To assess bulk BrdU incorporation levels as well as expose further BrdU epitopes, fixed cells were treated with DNase prior to immunostaining (data not shown). Briefly, individual coverslips were incubated with 10 units of RNase-Free DNase (M6101, Promega) in 1X Reaction Buffer (400mM Tris-HCl [pH 8.0 at 25°C], 100mM MgSO4, 10mM CaCl2) for 40 min at room temperature. After that, coverslips were washed once with Stop Buffer (20mM EGTA [pH 8.0 at 25°C]) in PBS and further washed two-three times with PBS. Coverslips were then equilibrated in Antibody diluent media for 15 min at room temperature before processing for immunofluorescence staining as described earlier.

### Quantification and Statistical analysis

Statistical analyses were performed on GraphPad Prism (Version 8.0). The details about the statistical tests are indicated in figure legends.

## ACKNOWLEDGEMENTS

We thank Laetitia Besse from the Multimodal Imaging Center Imaging Facility of the Institut Curie (CNRS UAR2016/ INSERM US43/ Institut Curie/ Université Paris-Saclay) for support with imaging and analyses. We thank the Institut Curie Next Generation Sequencing (Sylvain Baulande) platform for high-throughput sequencing. We also thank Aura Carreira and Martin Dutertre for helpful comments on the manuscript.

## AUTHOR CONTRIBUTIONS

M.L., C.B. and P.U. performed the CLIP, SIRF and 2C experiments. J.J. performed the gel shift and surface plasmon resonance experiments. G.C. and K.S. performed the QIBC experiments. P.B., D.B., X.V. performed recombinant protein production and associated molecular cloning. S.Me and S.Ma performed all other molecular cloning and laser micro-irradiation experiments. S.V., P.U., K.S, M.L. and C.B. conceived the experiments and analyzed data. S.V. coordinated studies and wrote the manuscript with the help of P.U., C.B., K.S., S.L. and the input from all other authors. Funding acquisition: S.V., K.S., P.B.

