## Supplemental Figure 1 for "53BP1 interacts with the RNA primer from Okazaki fragments to support their processing during unperturbed DNA replication"

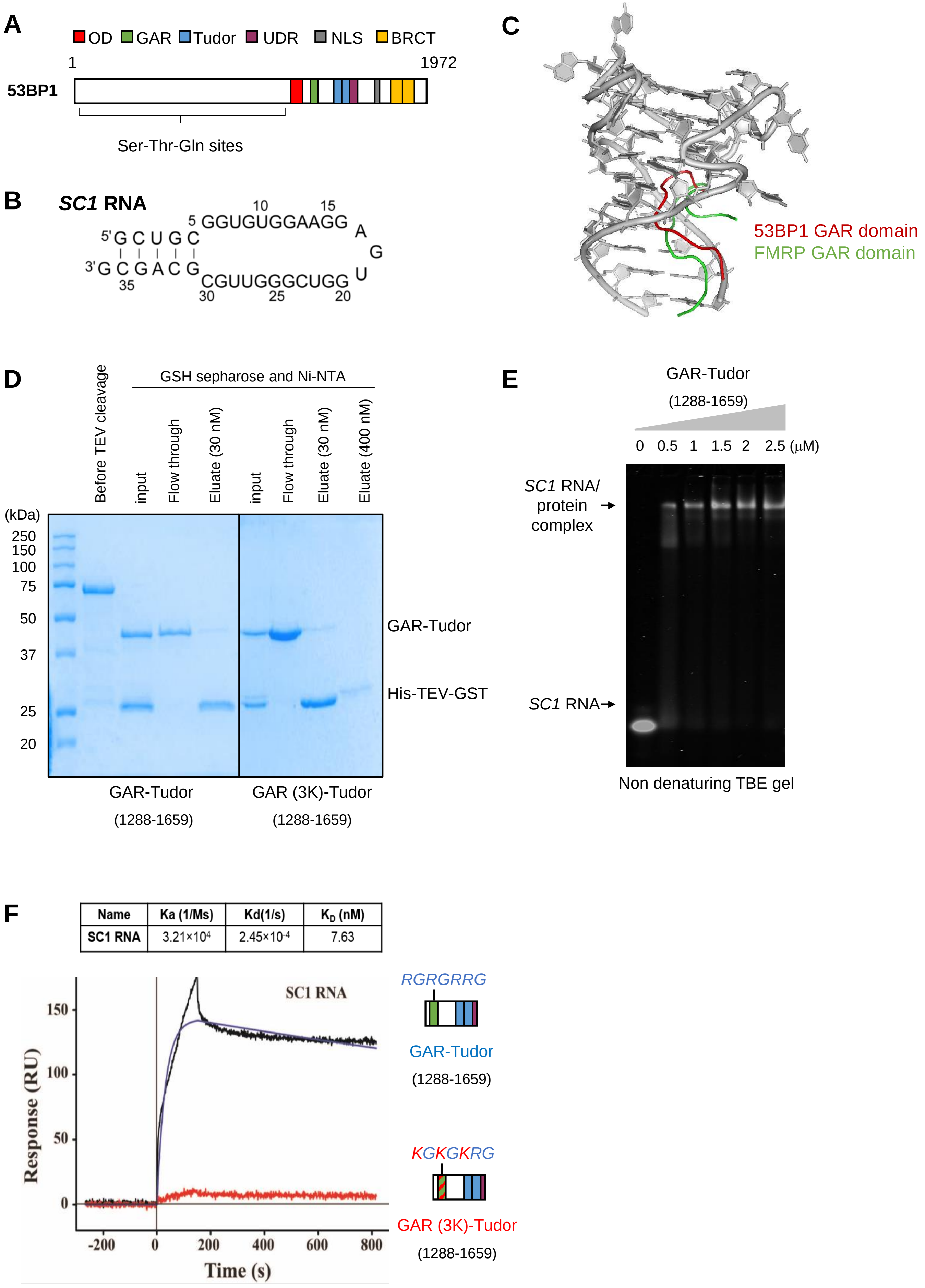


| Name | Ka (1/Ms) | Kd(1/s) | $K_D$ (nM) |
|---|---|---|---|
| SC1 RNA | $3.21\times10^{4}$ | $2.45\times10^{-4}$ | 7.63 |

**Supplemental Figure 1. 53BP1 binds RNA *in vitro.***
(A) 53BP1 domains map.
(B) Structure of *SC1* RNA.
(C) The predictive docked structure of *SC1* RNA (grey in colour) with GAR-TUDOR RGG motif (PRGRGRRGRP, red in colour) superimposed with X-ray crystal structure of the FMRP RGG peptide (GDGRRRGGGGRGQG, green in colour)–*SC1* RNA complex. The RGG motif of GAR-TUDOR and FMRP protein showed comparable binding mode with *SC1* RNA in the duplex–quadruplex junction.
(D) Gel purification of GAR-Tudor and GAR (3K)-Tudor recombinant proteins.
(E) Electrophoretic Mobility Shift Assay (EMSA) performed using purified recombinant 53BP1 containing the Tudor and the GAR domains incubated with a 5’-end fluorescently labelled synthetic 36 nt-long *SC1* RNA.
(F) Surface plasmon resonance (SPR) experiments with the indicated recombinant proteins and a 5’-end biotinylated *SC1* RNA.
