## Supplemental Figure 2 for "53BP1 interacts with the RNA primer from Okazaki fragments to support their processing during unperturbed DNA replication"

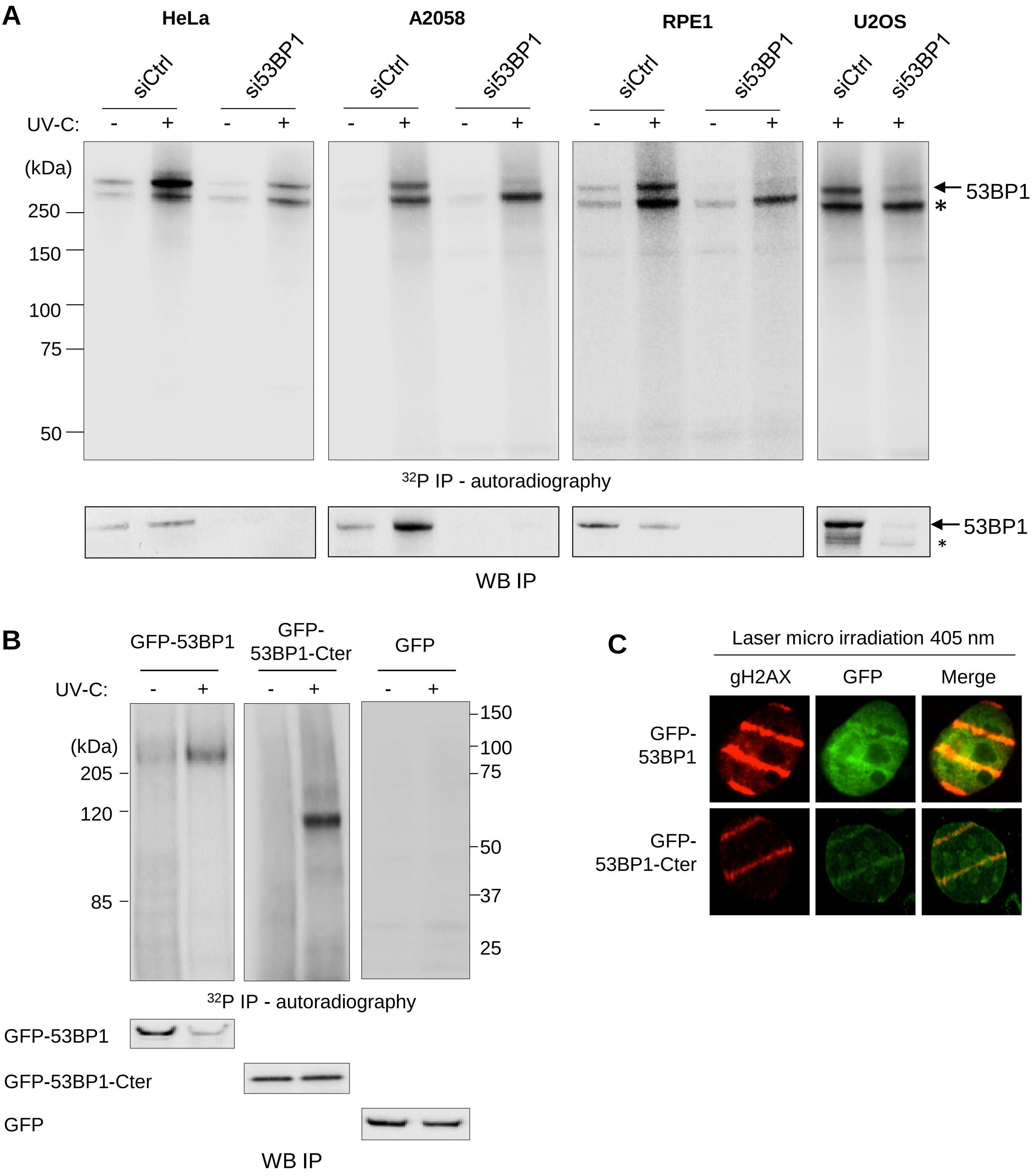

A
HeLa
A2058
RPE1
U2OS
siCtrl
si53BP1
UV-C:
(kDa)
250
150
100
75
50
53BP1
*
32P IP - autoradiography
WB IP
B
GFP-53BP1
GFP-53BP1-Cter
GFP
UV-C:
(kDa)
205
120
85
150
100
75
50
37
25
32P IP - autoradiography
GFP-53BP1
GFP-53BP1-Cter
GFP
WB IP
C
Laser micro irradiation 405 nm
gH2AX
GFP
Merge
GFP-53BP1
GFP-53BP1-Cter

**Supplemental Figure 2. 53BP1 directly interacts with RNA.**

(A) Autoradiography of 53BP1-nucleic acid complexes in UV-C treated (+) or untreated (-) cervix cancer cells (HeLa), melanoma cells (A2058), osteosarcoma cells (U2OS) or epithelial cells (RPE1) depleted of 53BP1 (si53BP1) or left untreated (siCtrl). The asterisk indicates a non-specific band (*), the arrow indicates the position of 53BP1-nucleic acid complexes.

(B) CLIP experiments performed in HEK29T cells transfected with the plasmids expressing the GFP-53BP1 proteins or the C-terminal part of 53BP1 (1235 – 1972 amino acids) or GFP alone. 10% of the IP was analysed by western blot to confirm immunoprecipitation (lower panel).

(C) Immunofluorescence was performed in U2OS cells expressing the GFP-53BP1 or GFP-53BP1-Cter (1235 – 1972 amino acids) proteins. The cells were irradiated with laser micro irradiation at 405 nm (stripes) and stained with anti- γH2AX (red) or GFP (green) antibodies.
