## Supplemental Figure 3 for "53BP1 interacts with the RNA primer from Okazaki fragments to support their processing during unperturbed DNA replication"

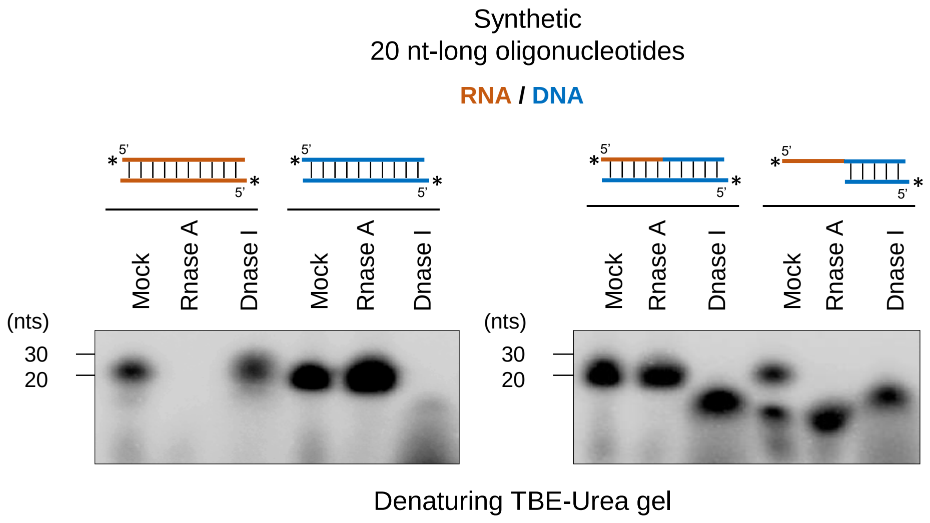


**Supplemental Figure 3. Enzymatic activities of RNAse A and DNAse I.**
*In vitro* assays of RNAse A and DNAse I activities on synthetic 20 nt-long oligonucleotides. The synthetic oligonucleotides were $^{32}$P-5'end labelled on both strands.
