## Supplemental Figure 4 for "53BP1 interacts with the RNA primer from Okazaki fragments to support their processing during unperturbed DNA replication"

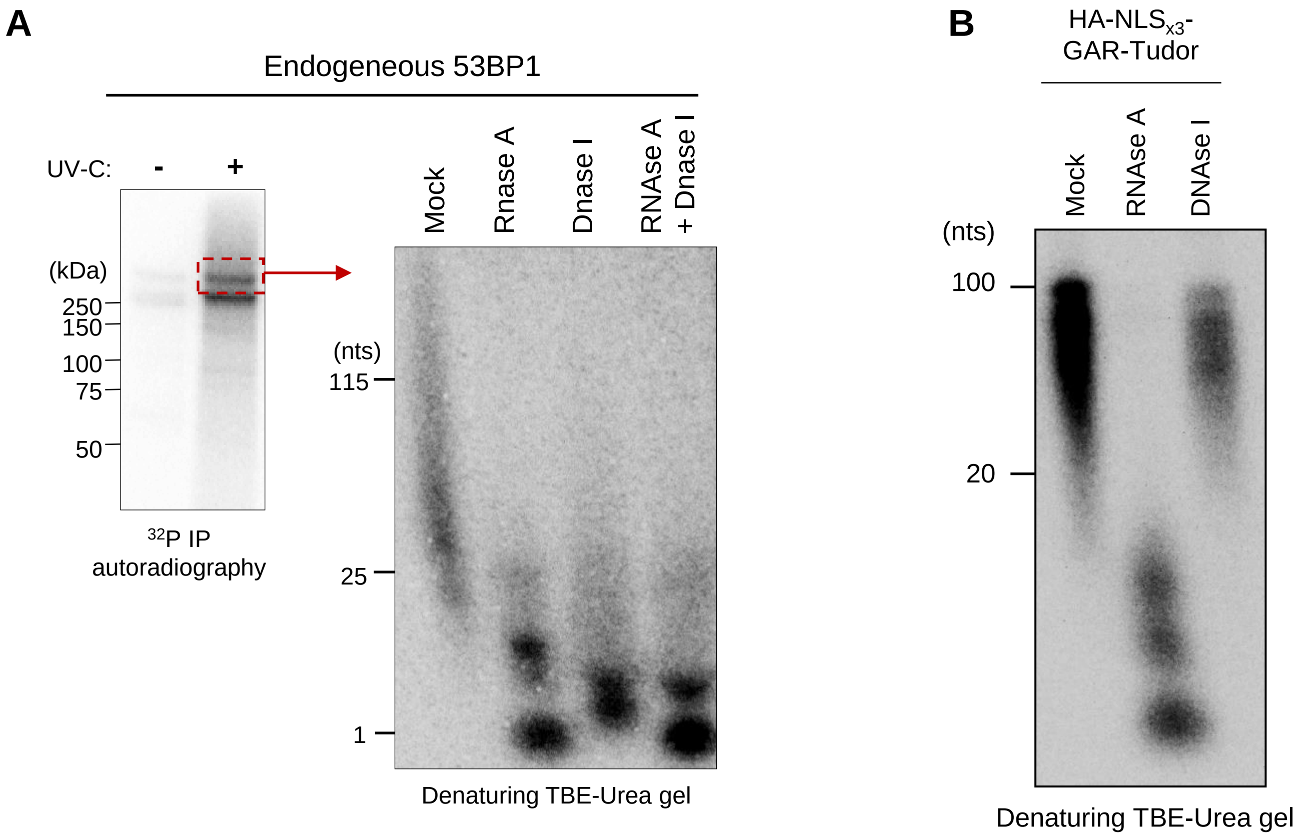


**Supplemental Figure 4. Endogenous 53BP1 binds RNA-DNA chimeras while the GAR-Tudor fragment does not.**
(A) Nucleic acid extraction after CLIP procedure of endogenous 53BP1 in A2058 cells.
(B) Nucleic acid extraction after CLIP procedure of HA-NLS$_{x3}$-GAR-Tudor fragment in transfected HEK293T cells.
