## Supplemental Figure 5 for "53BP1 interacts with the RNA primer from Okazaki fragments to support their processing during unperturbed DNA replication"

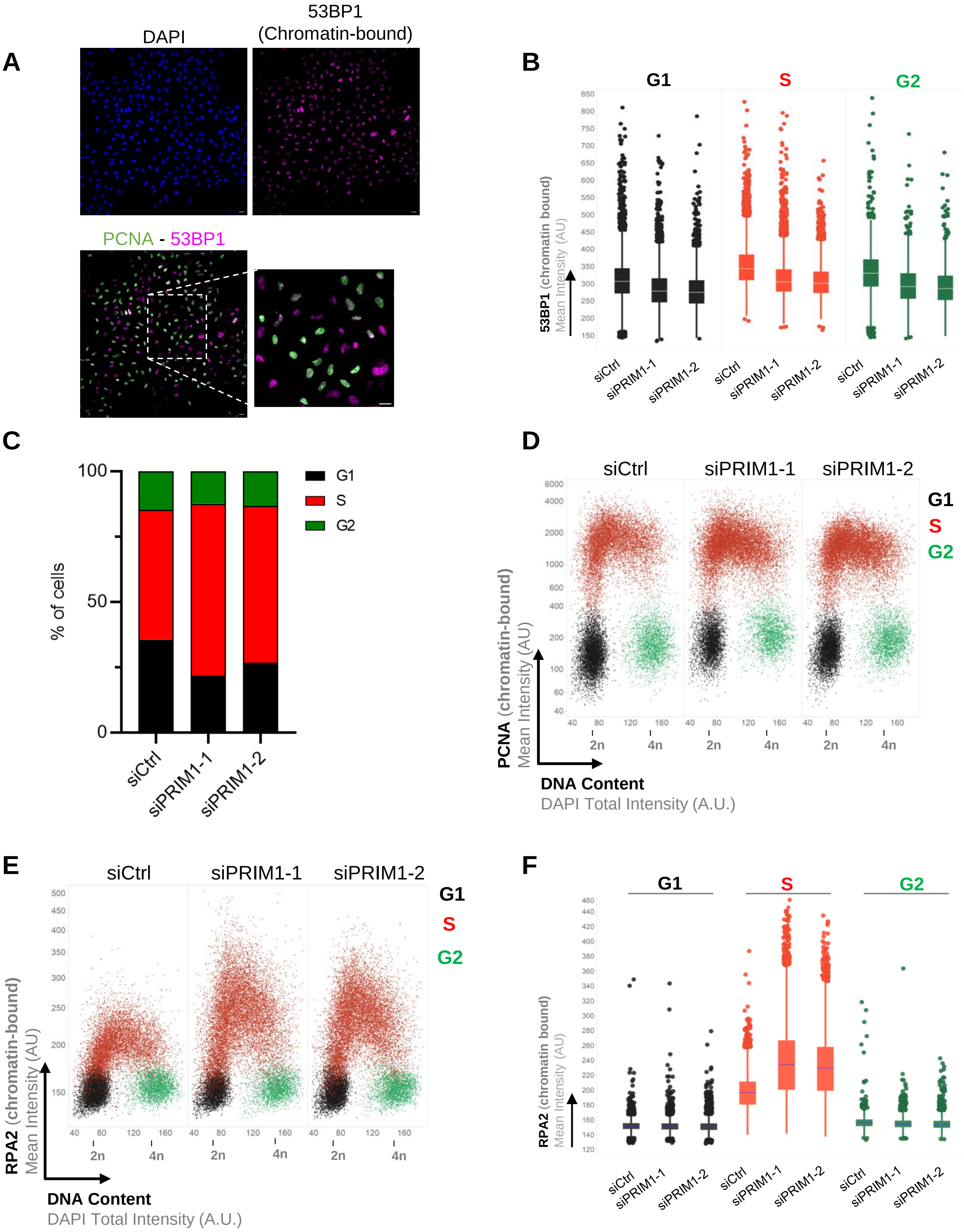
A
DAPI
53BP1 (Chromatin-bound)
PCNA - 53BP1
B
G1
S
G2
53BP1 (chromatin bound) Mean Intensity (AU)
siCtrl
siPRIM1-1
siPRIM1-2
C
% of cells
G1
S
G2
siCtrl
siPRIM1-1
siPRIM1-2
D
siCtrl
siPRIM1-1
siPRIM1-2
PCNA (chromatin-bound) Mean Intensity (AU)
DNA Content
DAPI Total Intensity (A.U.)
2n
4n
E
RPA2 (chromatin-bound) Mean Intensity (AU)
F
RPA2 (chromatin bound) Mean Intensity (AU)

**Supplemental Figure 5. Impact of PRIM1 depletion on the association of 53BP1, PCNA and RPA2 to chromatin.**

(A) Representative image of pan-chromatin and focus forming 53BP1 quantified in QIBC. Scale bar, 20 µm.

(B) QIBC-based quantification of chromatin loaded 53BP1 mean intensity in different phases of cell cycle (from experiment in Figure 3A). U2OS cells were treated with indicated siRNAs for 48 hours. In box plots, centre lines are medians, the boxes indicate the 25th and 75th centiles, the whiskers indicate Tukey values. P values were determined by one-way ANOVA with Tukey's test, **** $p<0.0001$ (n > 10,000 cells from combined all cell cycle stages per condition from 3 biological repeats).

(A) Percentages of cells from different cell cycle of the QIBC in (B)

(D) Cell cycle stages were gated based on PCNA mean intensity versus DAPI signals.

(E-F) QIBC of RPA2 chromatin loading in cells treated with indicated siRNAs for 48 h. Nuclear DNA was counterstained by DAPI. n > 10,000 cells per condition. In box plots, centre lines are medians, the boxes indicate the 25th and 75th centiles, the whiskers indicate Tukey values.
