## Supplemental Figure 6 for "53BP1 interacts with the RNA primer from Okazaki fragments to support their processing during unperturbed DNA replication"

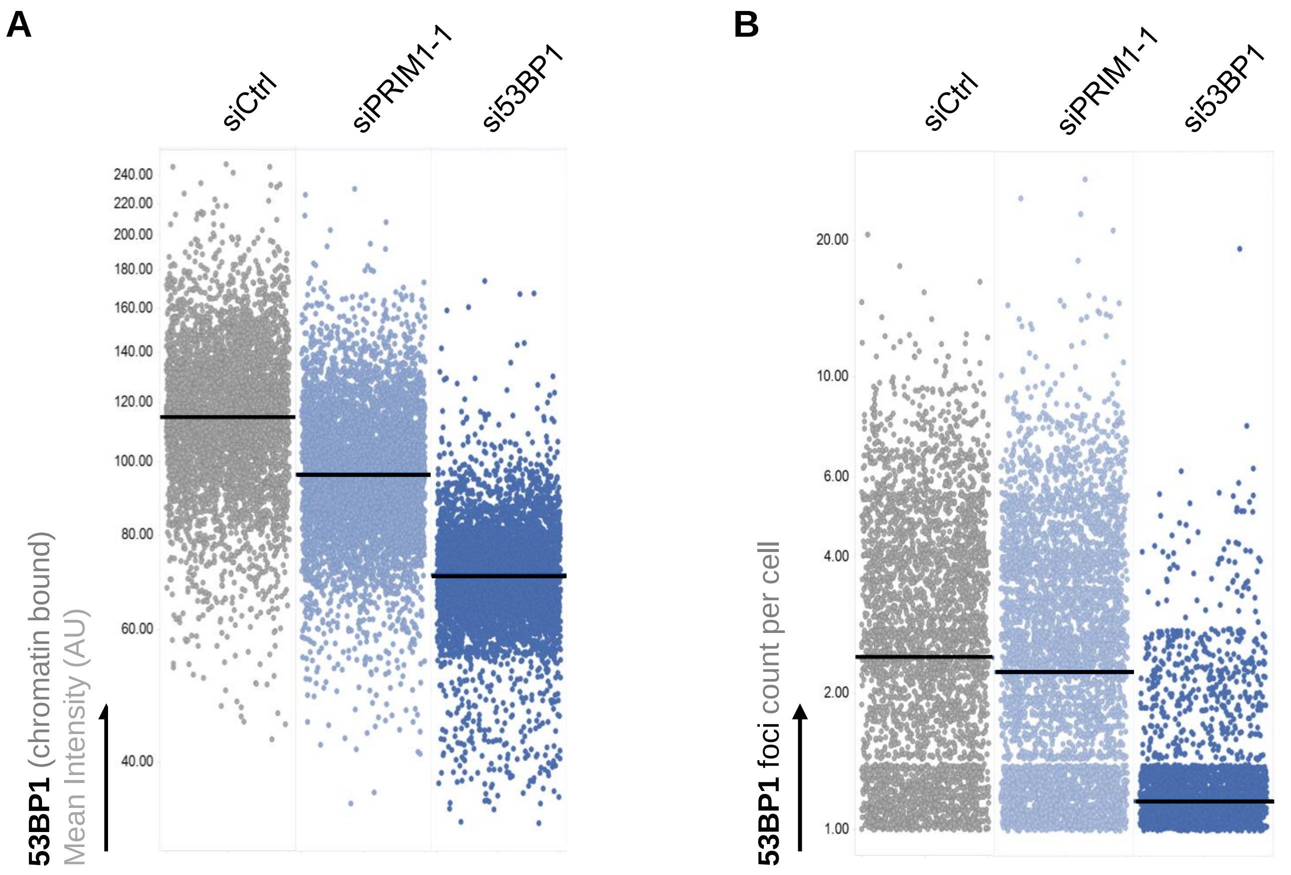


**Supplemental Figure 6. Loss of PRIM1 decreases 53BP1 at the chromatin without impacting the 53BP1 foci assembly.**

(A) QIBC of chromatin loaded 53BP1 mean intensity. Exponentially growing U2OS cells were treated with PRIM1 siRNAs for 48 h and 53BP1 siRNAs for 72 h. The horizontal lines are median values. n > 5,000 cells per condition from minimum of 2 biological repeats.

(B) Quantification of the 53BP1foci count derived from the QIBC analysis in (A). Average 53BP1 foci/cell are indicated with horizontal lines in black.
