## Supplemental Figure 7 for "53BP1 interacts with the RNA primer from Okazaki fragments to support their processing during unperturbed DNA replication"

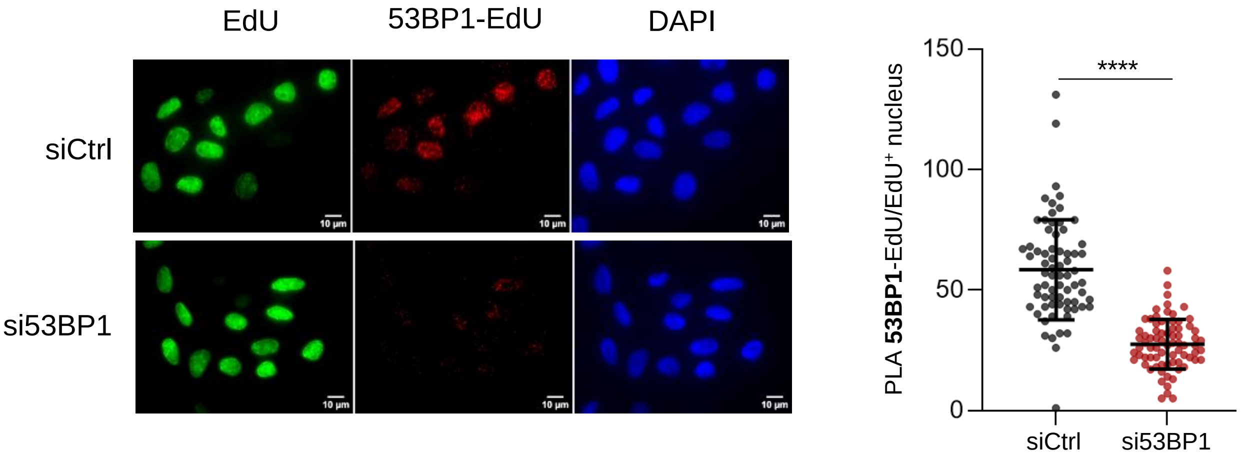


**Supplemental Figure 7. The PLA 53BP1-EdU signal in SIRF experiments is specific to 53BP1.**

Representative images of 53BP1-EdU SIRF experiments in U2OS cells transfected with indicated siCtrl or si53BP1.
