## Supplemental Figure 8 for "53BP1 interacts with the RNA primer from Okazaki fragments to support their processing during unperturbed DNA replication"

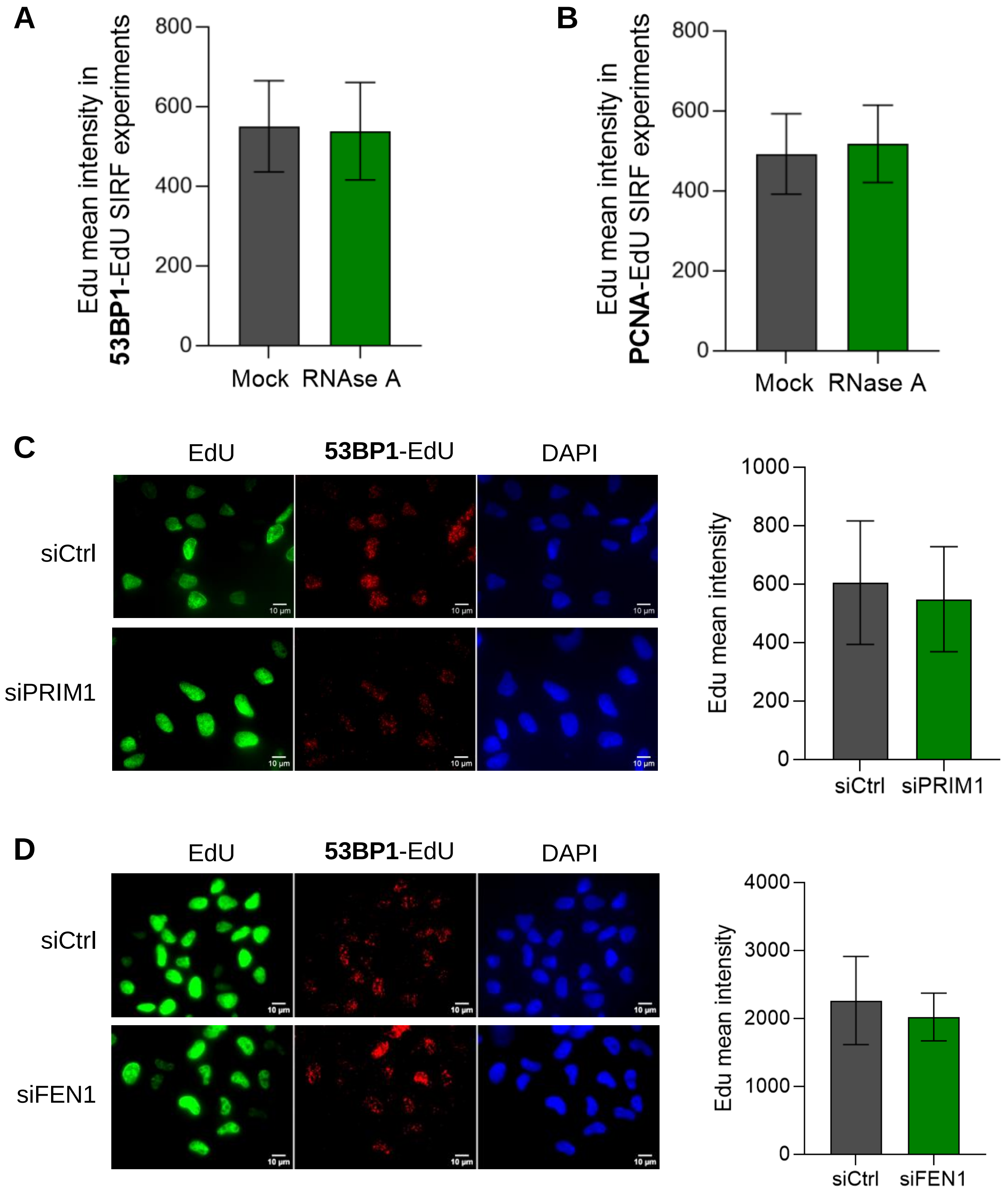


**Supplemental Figure 8. Intensity of EdU incorporation in SIRF experiments.**
(A-B) EdU mean intensity in 53BP1-EdU (A) or PCNA-EdU (B) SIRF experiments in U2OS cells treated or not with RNAse A.
(C-D) Representatives images of 53BP1-EdU SIRF experiments and EdU mean intensity in U2OS cells transfected with indicated siRNAs.
