## Supplemental Figure 9 for "53BP1 interacts with the RNA primer from Okazaki fragments to support their processing during unperturbed DNA replication"

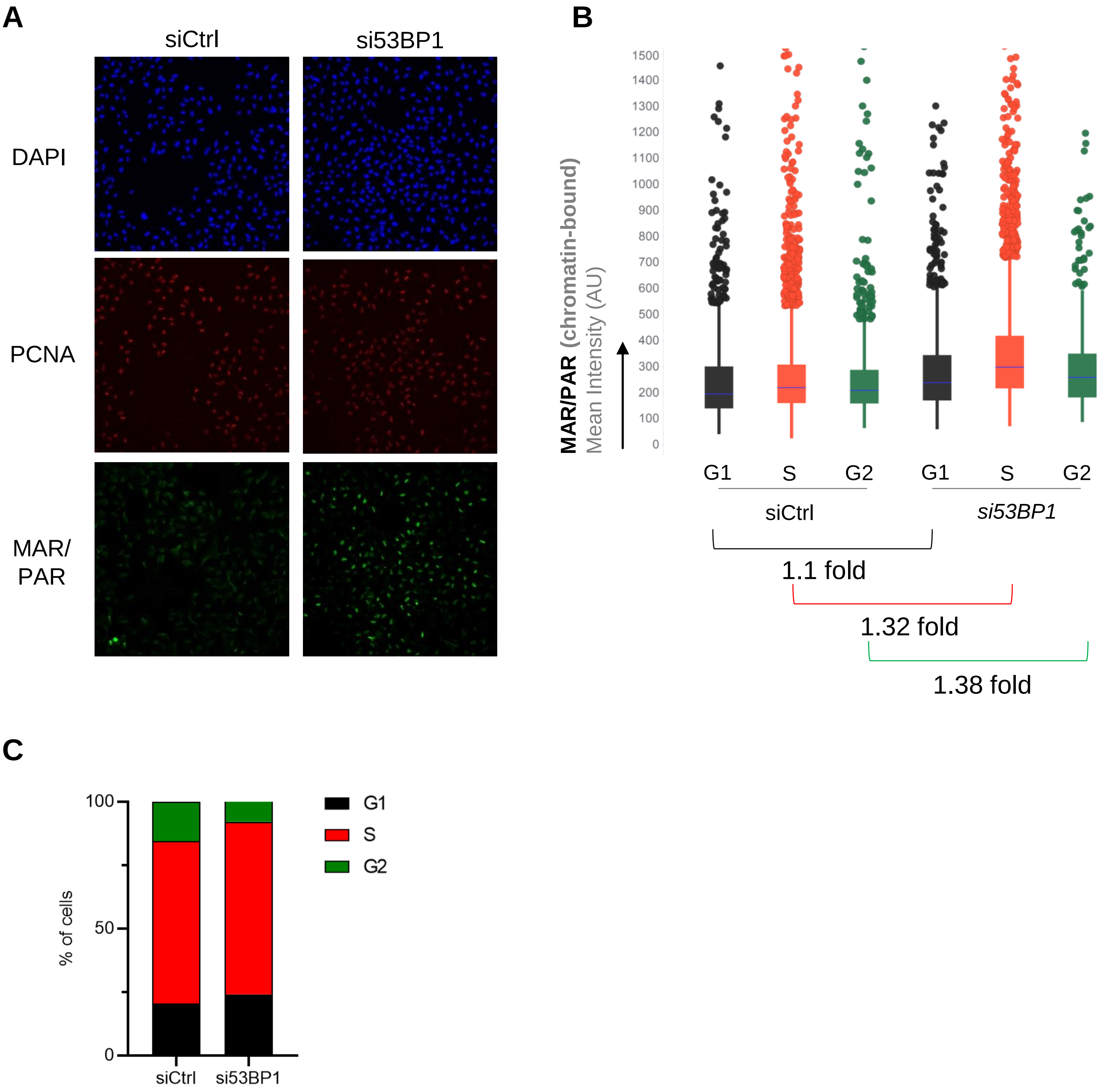


**Supplemental Figure 9. Images and quantifications of MAR/PAR bound to the chromatin.**

(A) Representative images of PCNA and MAR/PAR bound to the chromatin in QIBC experiment under si53BP1.

(B) QIBC-based quantifications of MAR/PAR signals from the experiment in A. In box plots, centre lines are medians, the boxes indicate the 25th and 75th centiles, the whiskers indicate Tukey values. P values were determined by one-way ANOVA with Tukey's test, **** $p<0.0001$ ($n > 10,000$ cells from combined all cell cycle stages per condition from 2 biological repeats).

(C) Percentages of cells from different cell cycle from the experiment in A and B.
