## Supplemental Figure 10 for "53BP1 interacts with the RNA primer from Okazaki fragments to support their processing during unperturbed DNA replication"

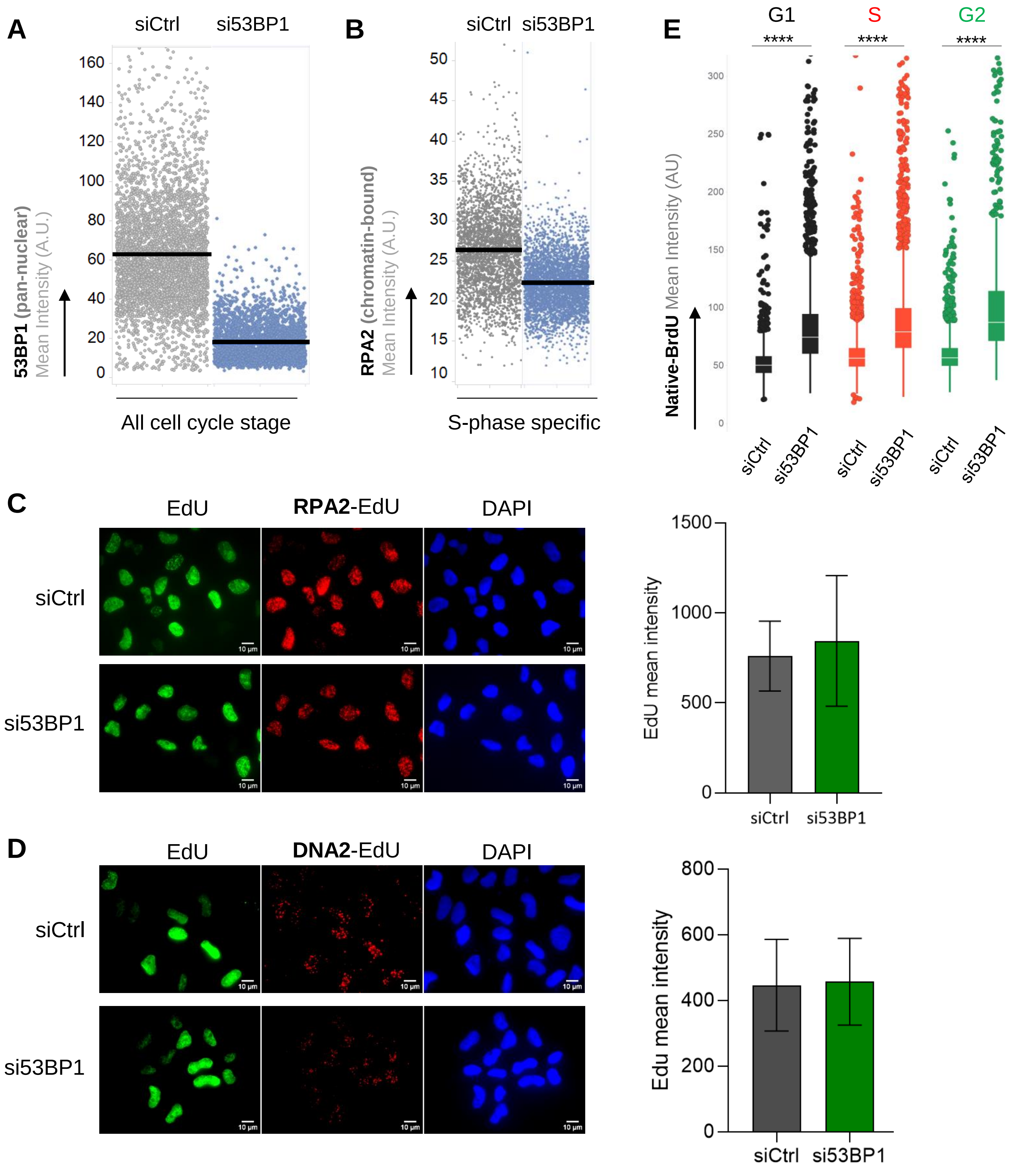

A
siCtrl
si53BP1
160
140
120
100
80
60
40
20
0
53BP1 (pan-nuclear) Mean Intensity (A.U.)
All cell cycle stage
B
siCtrl
si53BP1
50
45
40
35
30
25
20
15
10
RPA2 (chromatin-bound) Mean Intensity (A.U.)
S-phase specific
E
G1
S
G2
****
****
****
300
250
200
150
100
50
0
Native-BrdU Mean Intensity (AU)
siCtrl
si53BP1
siCtrl
si53BP1
siCtrl
si53BP1
C
EdU
RPA2-EdU
DAPI
siCtrl
si53BP1
1500
1000
500
0
EdU mean intensity
siCtrl
si53BP1
D
EdU
DNA2-EdU
DAPI
siCtrl
si53BP1
800
600
400
200
0
Edu mean intensity
siCtrl
si53BP1

**Supplemental Figure 10. Loss of 53BP1 decreases chromatin-associated RPA2 and increases ssDNA at nascent DNA.**

(A) QIBC of pan-nuclear 53BP1 mean intensity. Exponentially growing U2OS cells were treated with si53BP1 for 72h. The horizontal lines are median values. n > 5,000 cells per condition from minimum of 2 biological repeats.

(B) QIBC of chromatin loaded RPA2 mean intensity. Exponentially growing U2OS cells were treated with si53BP1 for 72h. The horizontal lines are median values.

(C-D) Representatives images of RPA2-EdU (C) or DNA2-EdU (D) SIRF experiments and EdU mean intensity in U2OS cells transfected with indicated siRNAs for 48 h.

(E) QIBC of ssDNA exposure upon depletion of 53BP1. Parental DNA in replicating U2OS cells was labeled by the addition of 10 μM BrdU for 24 h followed by a chase into normal medium for 1 h before cell fixing. Cells were fixed and stained with antibodies against BrdU without DNA denaturation to detect parental-strand ssDNA. Box plots of mean BrdU intensity per nucleus is shown. In box plots, centre lines are medians, the boxes indicate the 25th and 75th centiles, the whiskers indicate Tukey values. P values were determined by one-way ANOVA with Tukey's test, **** $p<0.0001$ (n > 5,000 cells from combined all cell cycle stages per condition from 2 biological repeats).
